# Assessing the degree of cortical dislamination through electrical pattern analysis

**DOI:** 10.1101/2025.06.16.659942

**Authors:** Ana Aquiles, Laura Pinedo-Vargas, Hiram Luna-Munguia, Fernando Peña-Ortega, Luis Concha, Román Rossi Pool

## Abstract

Focal cortical dysplasia (FCD) is a leading cause of pharmacoresistant epilepsy in pediatric populations, although its contribution to epileptogenesis remains incompletely understood. Recent findings indicate that hyperexcitability might stem from peripheral areas that are not dysplastic, rather than from the malformation itself. However, considering the significant variability associated with these malformations, it remains challenging to clarify whether this degree of disorganization contributes to changes in activity. In this study, we used the carmustine-induced animal model to investigate how varying degrees of cortical malformation influence neural dynamics. Local field potentials (LFP) were recorded using a multielectrode array (MEA) during both spontaneous activity and external perturbation. We developed a novel metric to quantify spatial heterogeneity in signal organization and evaluated its association with excitation-inhibition (E/I) balance. Our results reveal that alterations such as the aperiodic exponent value and the sparcity of clustering in signal classification are related to the extent and distribution of cortical abnormalities, underscoring the functional relevance of cytoarchitectural variability. This work advances the understanding of FCD-related network dysfunction and introduces analytical approaches with potential translational value for neuroscience research and pre-surgical evaluation.

## 1 Introduction

Focal cortical dysplasia (FCD) is the leading cause of surgical intervention for pharmacoresistant epilepsy in pediatric patients [1, 2]. The occurrence of FCD in cases of focal epilepsy is roughly 5-25 %. Yet, despite this prevalence, the relationship between FCD and epilepsy remains incompletely understood [1], as not every cortical malformation progresses to epilepsy [3]. It has been suggested that hyperexcitation in pediatric FCD type II may stem from the presence of balloon cells [4], which are primarily identified by their unique morphology. However, these balloon cells, despite their highly abnormal appearance, do not trigger epileptic activity, while other dysmorphic cells and immature neurons significantly contribute to the occurrence of epileptic discharges [5]. The combination of animal models and advanced analytical techniques has brought us closer to understanding the mechanisms of cortical dysplasias. Groundbreaking discoveries by Koh and collaborators [6], have challenged traditional views of cortical dysplasia, demonstrating that functional impairments may originate not within the primary malformation itself, but rather in surrounding non-malformed neurons. Their findings reveal that hyperexcitability often emerges from these ostensibly intact regions, while neurons within the malformed zones can remain hypoexcitable. The balance between excitation and inhibition is a fundamental concept to understand neural circuit deformation and brain dynamics alteration [7]. These principles have been increasingly explored to understand the emergence of epilepsy and other neurological conditions across various spatial and temporal scales [8, 9], including during the pre-ictal activity, which has been widely regarded as a critical window into the mechanisms leading to seizure onset [10, 11]. Yet, despite growing interest in describing and understanding early electrical changes in the development of epilepsy, the effect of cortical delamination caused by cortical dysplasia on functional alterations remains mostly unexplored. In this paper, we propose that baseline activity, particularly the local balance between excitation and inhibition, plays a key role in mediating the effect of varying malformation severity on neuronal dynamics. To address the latter, we employed the carmustine model to simulate the cytoarchitecture of cortical dysplasia [12–15], a model known for its wide variation in malformation severity, therefore closely reflecting the real challenge [16, 17]. Importantly, as this model does not present functional alterations as the evocation of seizure activity [17], we opted to measure spontaneous electrical activity using (MEA), and the Mg^2+^-Free model [18–23] to mimic the electrical signal evoked during epileptiform activity. Here, we decomposed the local field potential (LFP) spectrum to assess potential differences in signal structure based on cortical localization, and categorized those differences using frequency covariation. To quantify this categorization, we developed a metric that captures the heterogeneity of electrode groupings within each MEA recording. We then evaluated the aperiodic exponent in relation to an excitatory-inhibitory (E/I) balance metric [9, 24, 25], aiming to uncover a potential link with the degree of spatial sparsity in the categorization. Finally, we identified a common fluctuation band across different activity states (spontaneous, Mg^2+^-Free, and recovery), and tracked its temporal evolution relative to the initial degree of spatial categorization. Through this work, we aim to deepen the understanding of dysfunction in cortical dysplasia while accounting for the variability inherent in these malformations, while also offering analytical approaches that may inspire and support the study of complex biological time series in other systems characterized by structural and functional variability.

## 2 Results

### 2.1 Neural recordings from medial motor cortices during spontaneous activity, perturbation (Mg^2+^-Free) and recovery

To describe the electrical activity differences between healthy and cortical disarrangement tissues (i.e., Eulaminated and Dyslaminated, respectively), we evaluated M1 cortex in an animal model of cortical maldevelopment [17], incorporating a model that induces epileptogenic conditions. The Mg^2+^-Free model, previously reported and described [18, 23, 26, 27], was employed to assess how the absence of Mg^2+^ ions affects and how these changes relate to cortical disorganization. We compared signal differences in the medial motor cortex (see 4), where the MEA recorded activity spanning Layer II-III to V (Figure 2A). We first qualitatively examined the logarithmic power spectra (Figure 2B, bottom). Visual inspection of raw traces of untreated conditions (Figure 2C) across different states revealed noticeable differences in spontaneous and evoked activities (with Mg^2+^-Free alteration). It is evident from the spectrogams, even without formal analysis, that the primary distinctions between the conditions become apparent during spontaneous activity. Remarkably, these differences are much more pronounced during this resting period than during the other conditions. Nevertheless, it is also noteworthy that the appearance of both signals, evolved differently during the perturbation (Figure 2B bottom) challenge and continued during the recovery phase. The quantitative analysis of these observations is described in the following sections.

### 2.2 Categorization of electrodes and effects on the aperiodic exponent

Following the idea that a normal cortical lamination can lead to a functional structure, as shown by Dura-Bernal [28] based on real and simulated spikes train data (Figure 1), we first focused on description of spontaneous activity. We emphasized the categorization of electrodes within the MEA, using the distinct signal shapes associated with electrode depth (Figure 3A and Figure 1). We covary the frequency domain (60 points, from 0.5-100 Hz) over time for each electrode, and noted that the cumulative variance captured by the modes (components) 1 and 2 was greater in the Eulaminated condition than in the Dyslaminated one (Figure 3B). To further explore this result, we projected the first two modes on a comodulogram constructed from the spectrogram, ordered by electrode position in the MEA (Figure 3C), where we observed differences in frequency range and weighted magnitude across Modes 1 and 2. Mode 1 shows that the initial electrodes exhibit stable frequency and weighting values, while later electrodes display an upward trend in these values. Additionally, the Dyslaminated projection identifies several electrodes that display distinct behavior compared to the others. In module 2, variations in the electrode spectrum are also observed. This observation motivated us to look for groups in the space projection of the first two components. We obtained a variation of groups and dispersion along the MEA architecture, which led us to develop a metric to describe the degree of sparcity of the clusters. We refer to this metric as *ϵ*, (see 4), with values ranging from 0 to 1, where lower *ϵ* values indicate higher sparseness and less structured clustering, and higher values indicate the opposite (Figure 3D). Statistical analysis revealed significant differences in *ϵ* between the Eulaminated and Dyslaminated arrays, providing a measure to quantify the degree of disorder. Furthermore, based on previous results, these findings also highlight the range and variety of disorder, along with histological validation of the same model [16, 17]. Consequently, we suggest a theoretical link between the *ϵ* metric and cytoarchitecture, as illustrated in Figure 3D, bottom panel. As depicted in our initial observations (Figure 2), we next examined the aperiodic component found in each electrode of the MEAs to assess the observed variations. In Figure 3E, it is evident that the most extreme cases along the disorder gradient exhibit distinct power-law scale decay, based on the mean and variance of all measured electrodes. Interestingly, when evaluating the distribution of total aperiodic exponent values (Figure 3F), a bimodal function emerges, with a clear decline around 0.6 value. Based on this observation, we classified the aperiodic exponents using 0.6 as a threshold and re-evaluated the data using the same four representative examples shown in Figure 3D to investigate whether the previously identified clusters were preserved. This analysis revealed that Dyslaminated arrays predominantly exhibit aperiodic exponent values higher than 0.6, whereas Eulaminated arrays show a balanced distribution of values both below and above this threshold. To assess how well these values are related to the actual level of disorder described by *ϵ*, we performed a traditional linear regression between the bimodal coefficient of each individual array distribution and its *ϵ* value (Figure 4A). However, we found no significant correlation between them. Nevertheless, as shown in both Figure 4A and 4B, a tendency toward a bimodal distribution is apparent, where there exist both a high and low aperiodic values. Following the respective literature [24, 25], we propose a second theoretical approach to interpret this tendency : higher aperiodic exponent values could reflect an inhibitory predominance *I* ≫ *E*, whereas the lower values could correspond to an excitatory predominance *E* ≫ *I*. Despite the presence of electrodes that do not fit the previously mentioned categorization, it is evident that in Dyslaminated cortices, there is an imbalance between the two distributions. In contrast, in Eulaminated cortices, the highest and lowest values are evenly distributed in terms of the number of electrodes, which may play a significant role in the dynamics during an activity disturbance. We also conduct a comparison between groups similar to what is shown in Figure 3F, focusing on aperiodic exponential values during the subsequent activities, Mg^2+^-Free and Recovery. Both distributions appear to converge after an extended Mg^2+^-Free period, particularly during the 30-45 minute mark of perturbation. Additionally, during the recovery phase, the Eulaminated and Dyslaminated groups continue to exhibit a unimodal distribution, with the peak observed around the 1.0–1.2 range. This likely represents the most stable aperiodic value, as it persisted from the baseline activity.

**Figure 1:**
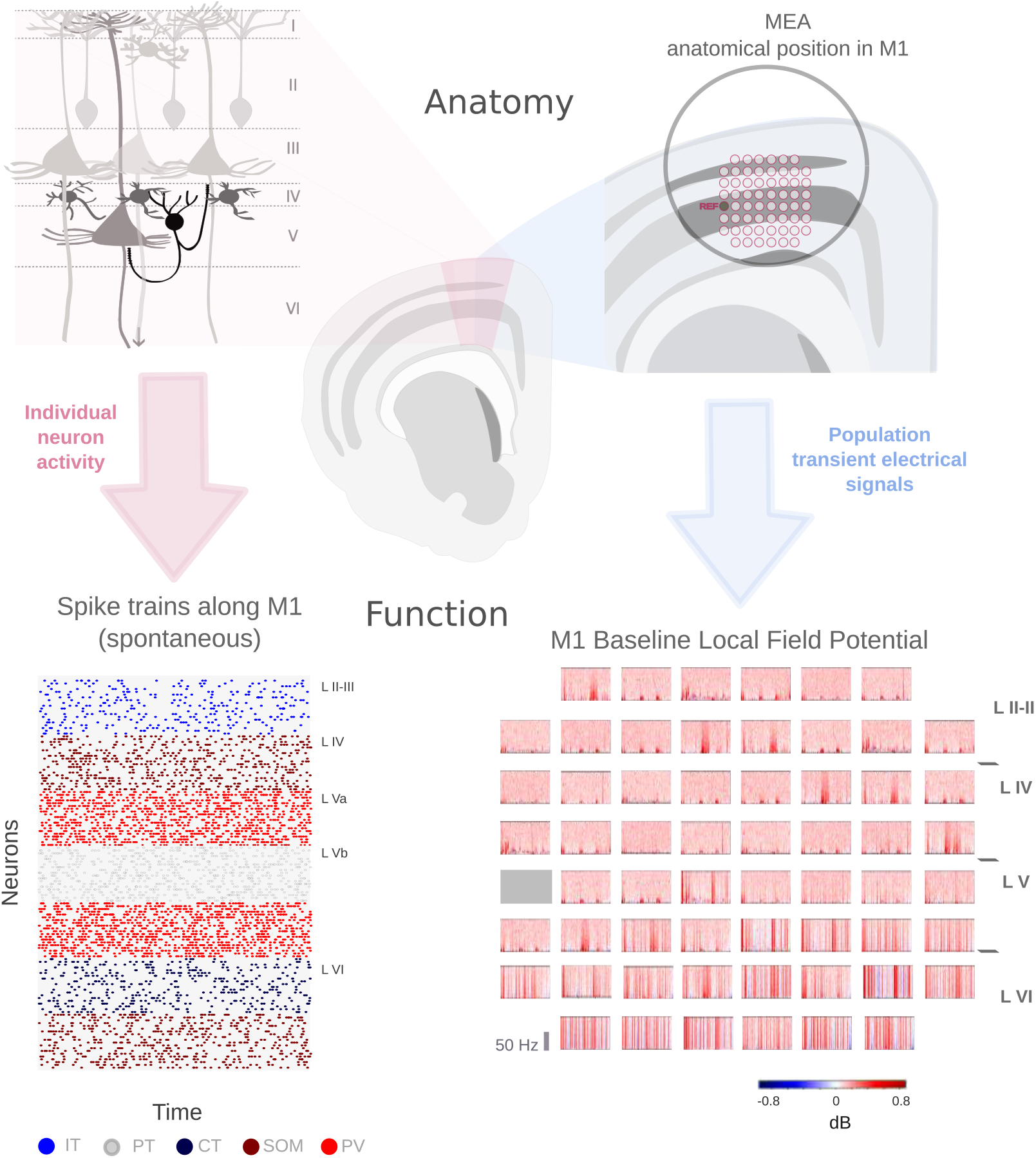
Hypothesis. Taking a similar approach to that of Dura-Bernal et al. (2023), who classified neuronal types based on neuronal firing patterns, we propose categorizing electrodes by their activity using temporal frequency covariations, as measured by LFP signals.

**Figure 2:**
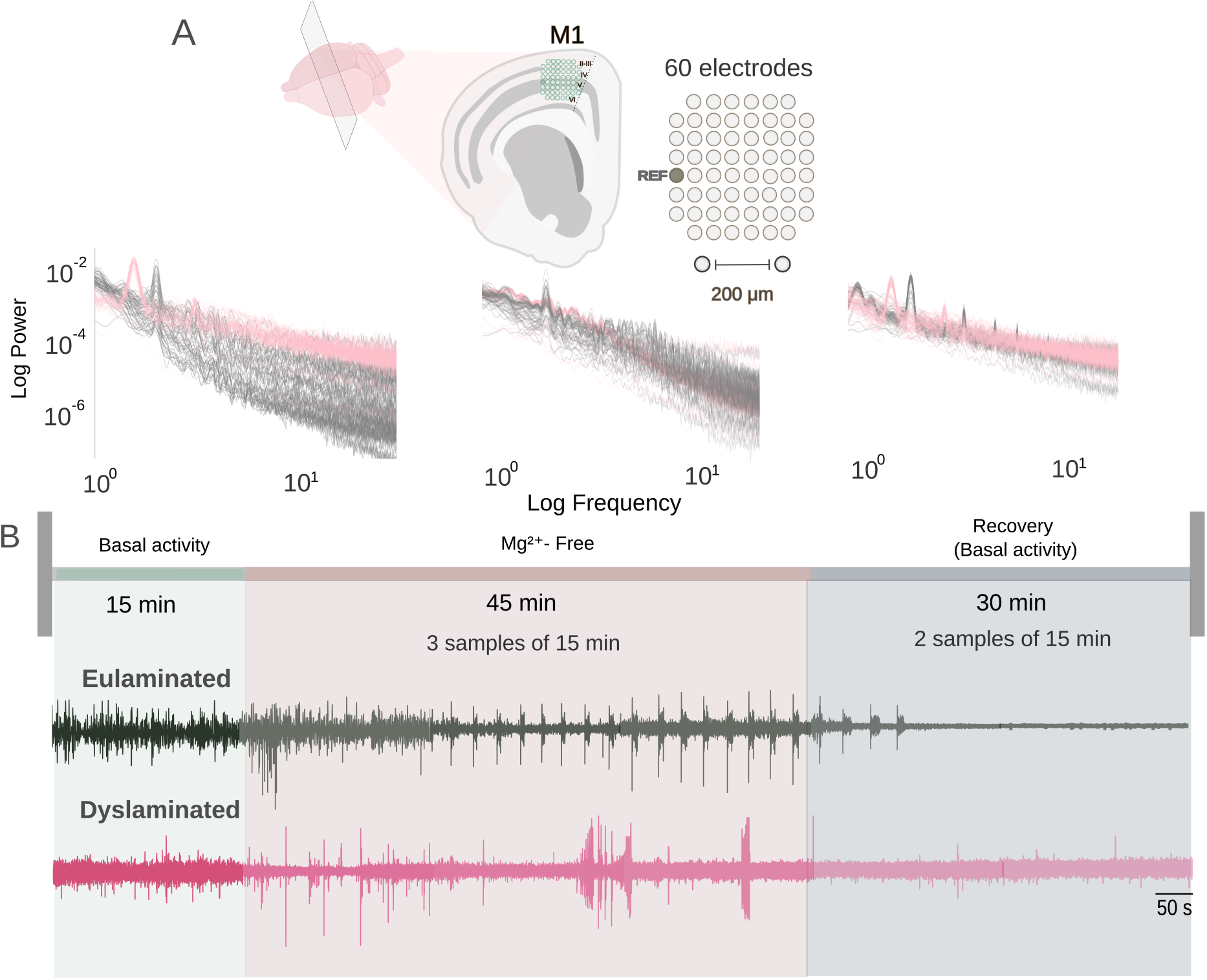
Overview. Multielectrode array recordings from normal (Eulaminated) and Dyslaminated cortexes. A) Approximate anatomical position of recordings, and spectral decomposition of spontaneous activity under three conditions in each of the 60 electrodes, for one control (grey) and one animal with dyslaminated cortex (pink). B) Exemplary time courses of spontaneous cortical activity under the three conditions. Each sample was about 15, 45 and 30 min per activity type (green - Basal activity, pale rose - Mg^2+^-Free, pale violet - Recovery). Examples were selected from the extreme cases of each condition.See also Figure 3

**Figure 3:**
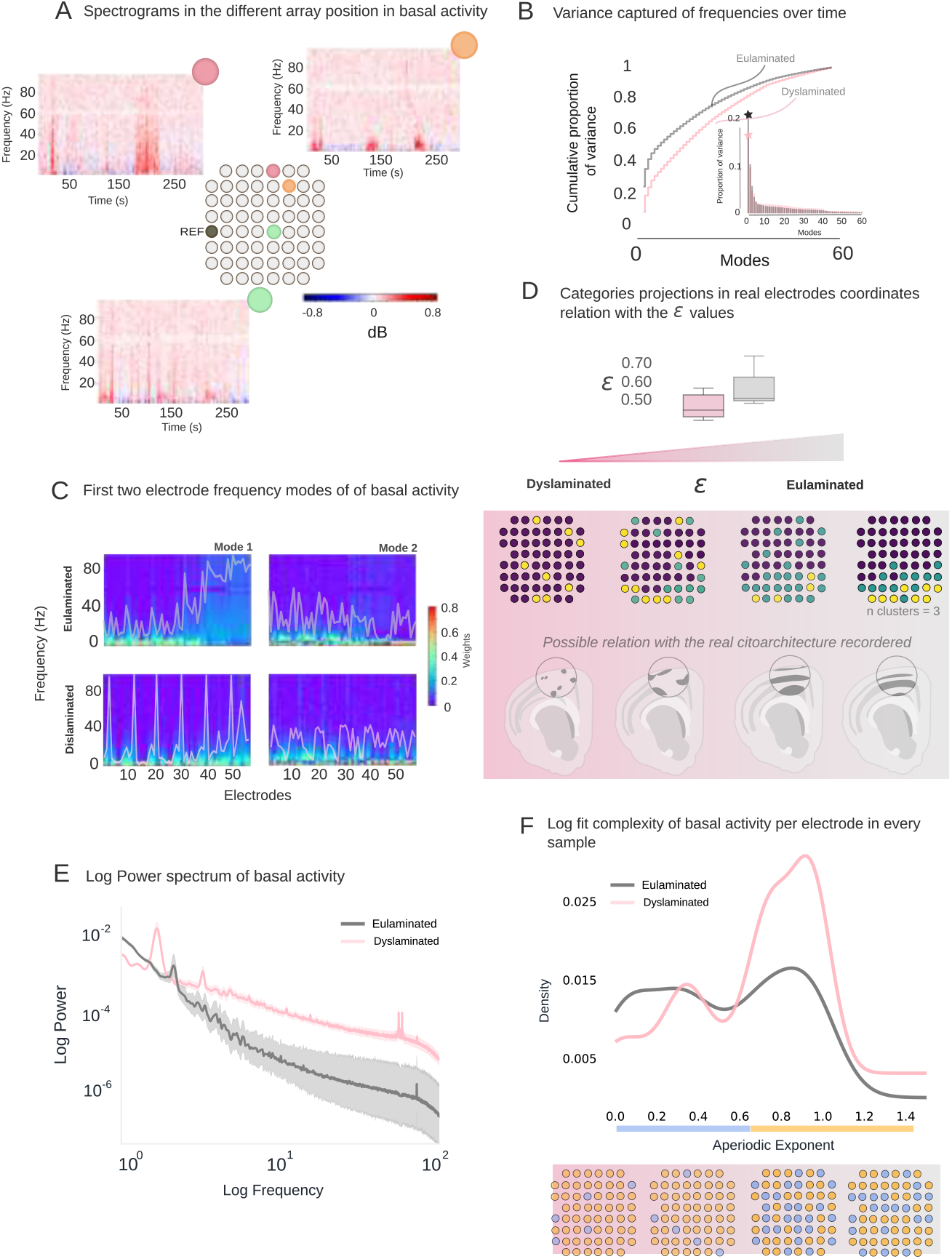
Classification of electrodes based on baseline activity to differentiate between Eulaminated and Dyslaminated cortexes. Temporal frequency covariation. A) Normalized spectrograms showing baseline activity across various electrode positions. Example shown corresponds to a Eulaminated extreme (*ϵ* = 0.75). Additional Dyslaminated and Eulaminated examples of complete spectrogram arrays can be found in Supplementary Figure 2. B) Variance captured from the frequency covariance in both groups (extreme examples, *ϵ* = 0.75 and *ϵ* = 0.26); Eulaminated depicted in light grey, Dyslaminated in light pink. C) Projection of the first two principal components (modes) into the comodulogram of frequency variation across all electrodes. The color scale indicates PCA weights; white lines illustrate mode fluctuations across electrodes. D) *ϵ* values from the layered disarray metric show a significant difference (p ¡ 0.01). Electrode categories (n=3), i.e. projection of the first 2 modes into real electrode positions, potentially linking to the level of cortical disorganization (hypothesis), at bottom. The extremes correspond to the same *ϵ* values as in previous panels. E) Logarithmic power spectrum for each electrode (extreme cases) along with aperiodic exponent values fitted across all records analyzed (Eulaminated = 4; Dyslaminated = 5) F) Distribution of aperiodic exponent values. See further details in Supplementary Figure 1 and 2.

**Figure 4:**
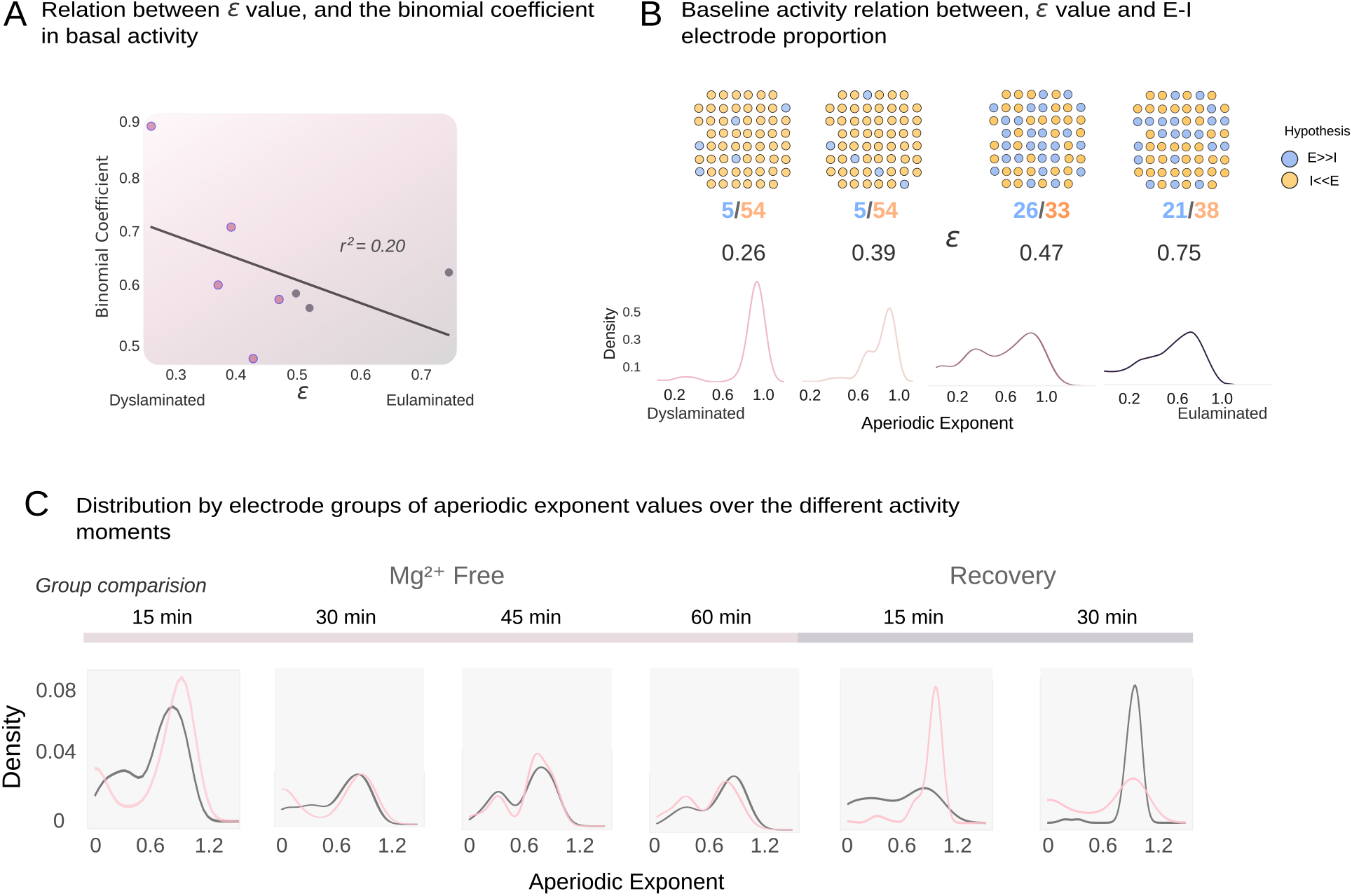
Relationship between the aperiodic exponent and the gradient of disorder in baseline activity, and its changes during perturbation and recovery. A) Linear regression between bimodal coefficient and *ϵ* values, *r*^2^= 0.20, p ¿ 0.05. Pink dots represent Dyslaminated arrays; grey dots represent Eulaminated ones. B) Electrode projection for various instances of disorder, with corresponding *ϵ* values displayed at the bottom. Percentages of aperiodic exponent values above and below 0.6 highlighted in orange and blue, respectively and the hypothesis that values under 0.6 signify heightened excitation *E* ≫ *I*, while those above indicate increased inhibition *I* ≫ *E*. This establishes a connection between the aperiodic exponent and the degree of disorder in baseline activity, as well as its changes during perturbation and recovery. The specific arrangement of aperiodic values for the selected examples is depicted in the lower panel. C) Comparison of the distribution of aperiodic exponent values across groups during consecutive activities: perturbation (Mg^2+^-Free), and recovery.

### 2.3 Functional changes in typical bands during perturbation and recovery

To assess functional differences associated with perturbation induced by a Mg^2+^-Free medium and the subsequent recovery period, we analyzed a shared range of frequency fluctuations. This choice was motivated by established evidence that human seizure discharges often exhibit coupling between low and high frequencies [10, 29]. Our goal was to identify the frequency range showing the greatest variation in our recordings, drawing inspiration from Zhang, W.[30], who reported a consistent fluctuation band by analyzing temporal covariation in the power spectrum. We adapted their approach, considering that no specific frequency band has been universally defined for evoked epileptic activity. Our analysis revealed that the highest covariation occurred in the lower frequency range (0.5–30 Hz), and this effect was more pronounced in Dyslaminated cortices. We demonstrated this through the projection of the first two components of spectral covariation onto a weighted spectrogram and by projecting both modes onto a weighted power spectrum (Figure 5A., B). Notably, covariance during baseline was consistently greater in Eulaminated cortices than in Dyslaminated ones (Figure 5C, upper panel). In contrast, during the Mg^2+^-Free perturbation, cumulative variance between the two groups converged (Figure 5C, lower panel). This pattern is further illustrated in the representative logarithmic power spectra (Figure 5B, right), where baseline bandwidths maintain divergent decay slopes, while the Mg^2+^-Free condition shows spectral convergence. This confirms our initial qualitative observations from Figure 2B, now filtered for the frequencies showing the highest covariance. To validate that the outcome of this bandwidth investigation preserves the ripple activity traits of the Mg^2+^-Free model, we mapped the bandwidth threshold onto three electrodes chosen at random from different deepest MEA positions (upper, middle, and lower). As illustrated in Figure 5D, the observed activity changes persist under this constraint. To track the evolution of activity in relation to the previously defined levels of disorder, we analyze the transfer of information between the electrode categories identified, using the common band variation, (see Figure 3). Starting with baseline recordings (Figure 6A), we clarify that Dyslaminated samples conveyed lower information values compared to Eluaminated ones. Additionally, where the MEA shows lower *ϵ* values, the transfer of information becomes less structured and primarily occurs from the largest electrode category (purple dots) to the sparsity one (yellow dots).

**Figure 5:**
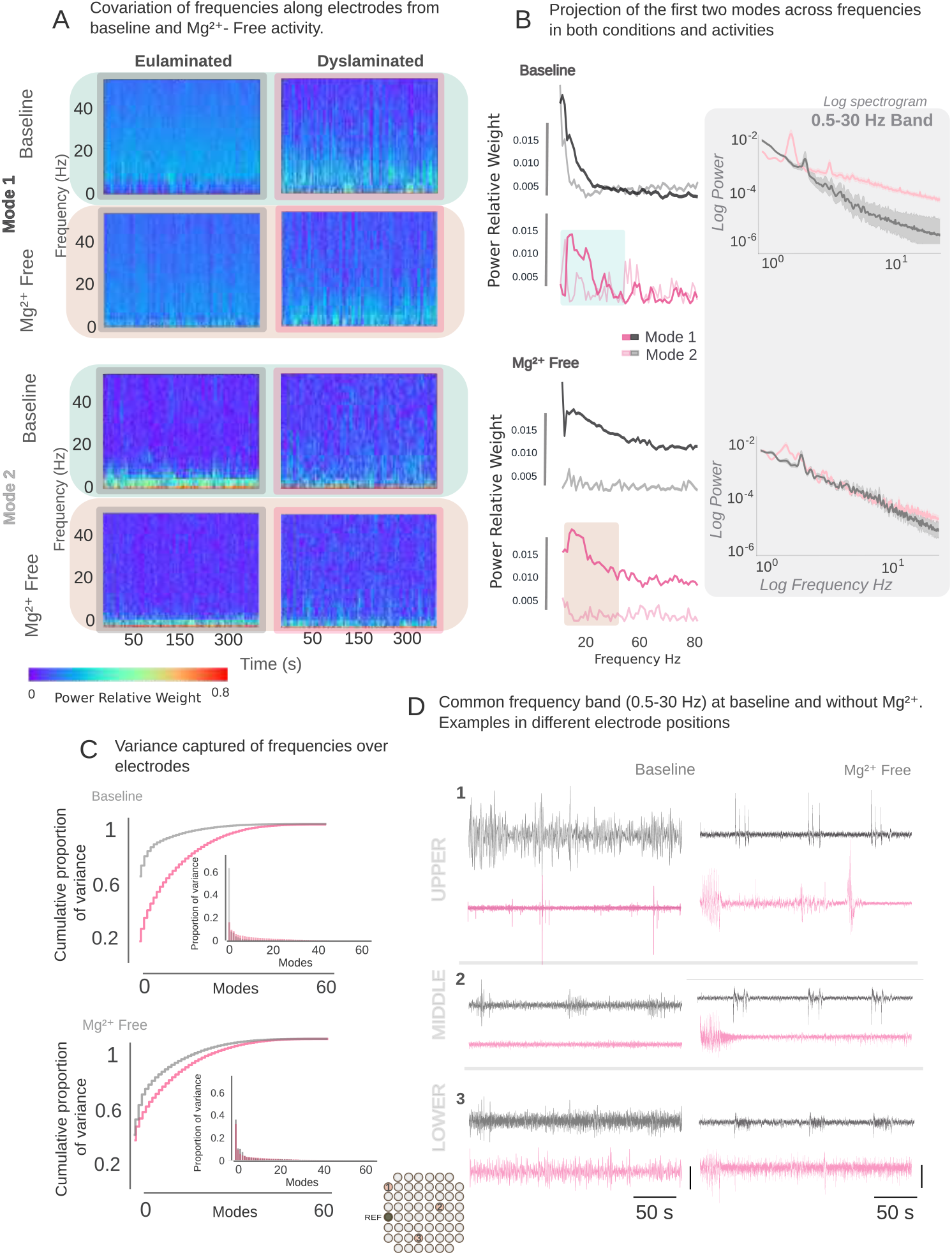
Common frequency band (0.5-30 Hz) selection from Dyslaminated cortices. Electrode frequency covariation. A) Spectrograms adjusted to highlight the first two modes during basal activity (depicted in green) and in Mg^2+^-Free (shown in pale rose). Examples from extreme conditions. See Figure 2. B) The power spectrum weighted according to the first two modes averaged across all samples (Eulaminated = 4; Dyslaminated = 5). The shaded region indicates the frequency range where the power was most prominent. C) Variance distributions captured in both conditions (baseline and in Mg^2+^-Free). Light pink corresponds to Dyslaminated cortices and light grey to Eulaminated. D) Examples of baseline and Mg^2+^-Free activity after filtering by the common frequency band, shown across three MEA depths (lower, middle, upper). Examples in panels C and D correspond to the extreme conditions. See further details, in Supplementary Figure 3.

**Figure 6:**
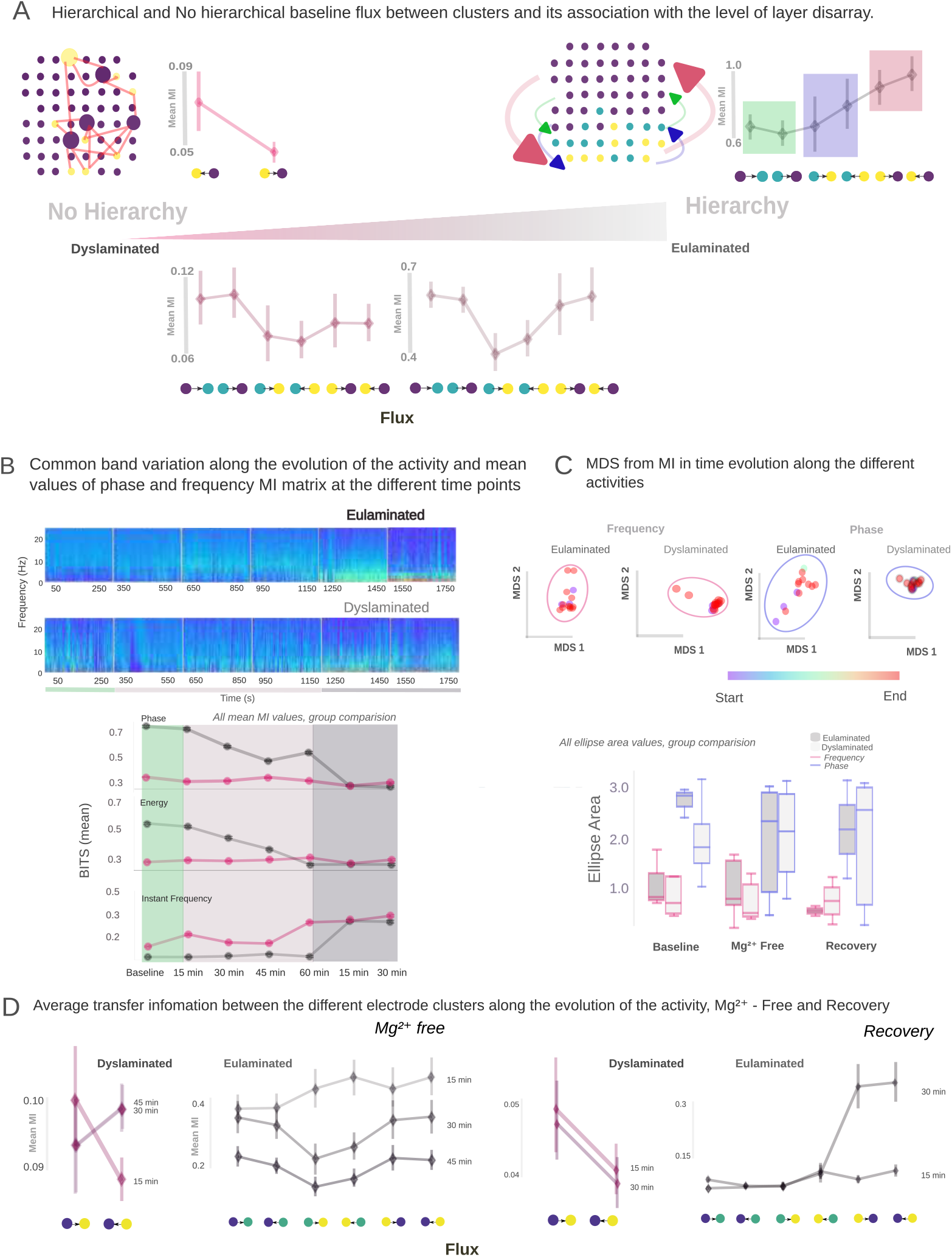
Global dynamical evolution of the common frequency band and its relationship with information flow across electrode categories. A) Baseline information flux for extreme Dyslaminated and Eulaminated samples, along with two intermediate cases (*ϵ* = 0.26, *ϵ* = 0.75, respectively). B) Average spectrogram within the shared band defined in Figure 3, representing total activity captured across conditions (Baseline in light green, Mg^2+^-Free in pale pink, Recovery in pale violet). Bottom: mean information (in bits) over time for each signal component within the common band, Phase, Energy and instantaneous frequency. C) MDS analysis of dissimilarity in mutual information matrices for each activity condition, with ellipses fitted at each point to represent the spread of dynamics in low-dimensional space. Right: temporal evolution of frequency; left: temporal evolution of phase. Bottom: ellipse area values across all recordings and activity conditions. D) Information flux during Mg^2+^-Free and Recovery phases for the same extreme dyslaminated and eulaminate samples shown in panel A.

It appears that the sparsity category transmits less information compared to the other, which may correspond to the hyper and hypoexcitable regions previously suggested by Koh, H .[6] but this time interpreted in terms of information content. However, as the *ϵ* values increased, a hierarchical pattern began to emerge in the flow of information, progressing from deeper electrode categories to more superficial ones. This observation seems to align with the known hierarchical organization of the motor cortical circuit [31], particularly in the zone where we attempted to record Figure 2A.

We next asked which component of the signal carries the highest amount of information. Following the approach of Kayser, C.[32], where they found that information amounts are distributed differently in the distinct components of a LFP signal, we analyzed the mean weighted spectrogram across the three activity paradigms (Baseline/Spontaeous activity, Mg^2+^-Free and Recovery) and evaluated the information content of three signal components: phase, energy, and instantaneous frequency (Figure 6B, lower panel). These values reflect the average trend across all MEAs within the common LFP bandwidth, without distinguishing between levels of delamination, as the observed variability was insufficient for clear separation. Our results show that phase carries the highest amount of information across conditions, while instantaneous frequency contributes the least. Furthermore, phase and energy follow similar trends over time. In Eulaminate cortices, both components begin with high information values during baseline activity but decrease during Mg^2+^-Free perturbation and do not return to baseline levels during recovery. In contrast, Dyslaminated cortices also begin with high values, but their information content remains more stable over time. Additionally, the trends of phase and energy appear to evolve similarly over time in both conditions; on one hand, Eulaminated phases and energy begin with high information values, but as perturbation continues, they decrease and never return to their original levels, even during the recovery state. On the other hand, the Dyslaminated values also initiate at high information levels, but their trend shows minimal change over time. However, instant frequency shows the least informative values in both situations. Additionally, the pattern seen suggests that the Dyslaminated cortices seem to enhance their information content as time goes on and are likely to show higher values compared to the Eulaminated, during spontaneous activity and in the absence of Mg, but again appear similar in the recovery phase, as with the other two components.

### 2.4 Temporal evolution of signal components

To better understand the temporal evolution of these components, we focused on the most informative component, the phase, and the least informative, the instantaneous frequency. Using mutual information computed between electrodes across the full shared LFP bandwidth and a sliding window approach, we quantified the dissimilarity between each instantaneous state and its subsequent state within each window (see 4). Dimensionality reduction was then applied to these temporal dissimilarity matrices to track the evolution of each component across different activity paradigms (Figure 6C). The upper panel presents a representative example from the extremes of the *ϵ* distribution. Each point corresponds to a time window, with color indicating temporal progression (red = early, blue = late). The ellipses represent the low-dimensional projections of the dynamics: red for frequency and blue for phase. For each multidimensional scaling (MDS) projection, an ellipse was fitted to quantify the spread of the dynamics; larger ellipses indicate greater variability and a higher capacity to transform the underlying interaction dynamics. The boxplots beneath these projections reveal that frequency dynamics were more temporally constrained, with slightly higher mean values observed in eulaminate cortices than in dyslaminated ones, except during the recovery phase. This observation aligns with the reduced information content shown in Figure 6B, supporting the interpretation that lower information corresponds to a diminished capacity for dynamic change. In contrast, phase dynamics exhibited greater variability in dyslaminated cortices across all activity paradigms, whereas eulaminate cortices showed increased variability only during the Mg^2+^-Free condition. This highlights a key role for phase in modulating neural interactions under perturbed conditions. As with frequency, the average ellipse sizes for phase were generally larger in eulaminate cortices, except during recovery, where dyslaminated cortices showed increased variability that corresponded to elevated phase information content (Figure 6B). These findings offer valuable insights for future analytical decompositions and are consistent with previous studies emphasizing the importance of phase locking across frequency bands [11, 32]. Building upon the prior decomposition analysis, we re-examined the information transfer depicted in Figure 6A, this time focusing on distinct phases of activity while maintaining a consistent bandwidth. Figure 6D illustrates the same extreme cases from Figure 6A, now under Mg^2+^-Free conditions (left) and during the Recovery phase (right). In dyslaminated cortices, average information transfer shows a slight increase; notably, during the first 15 minutes of Mg^2+^-Free exposure, the transfer flux remains aligned with baseline levels. However, this trend reverses over the subsequent 30 minutes. In contrast, eulaminate cortical activity displays a progressive decline in information transfer relative to spontaneous activity, with an early disruption of hierarchical organization evident from the onset of Mg^2+^-Free exposure. During recovery, the dyslaminated cortex rapidly re-establishes its original transfer pattern within the first 15 minutes, whereas the eulaminate cortex requires 30 minutes to partially restore hierarchical flow and fails to fully recover it. Collectively, these findings suggest that a certain degree of structural disorganization may impart functional resilience to external perturbations. This disorganization appears to affect not only the excitatory–inhibitory balance, but also the overall efficiency and stability of information transmission within the network. These insights may have broader implications for understanding how cortical circuits respond to and recover from functional disruptions, and how structural variability modulates their adaptive capacity.

## 3 Discussion

Surgical resection remains the primary treatment for cortical dysplasia, largely due to its strong association with pharmacoresistant epilepsy, particularly in pediatric populations [1, 2]. This significant clinical challenge motivated our continued investigation of the carmustine model during early stages of neurodevelopment. Although this model presents certain limitations, such as its embryonic onset and the production of regionally diffuse and heterogeneous malformations [12]; we viewed these features as opportunities rather than constraints. By leveraging advanced analytical tools, we sought to characterize the extent of cortical disruption and assess its functional consequences, particularly in response to external perturbations. Focusing on spontaneous activity, we classified electrode behavior to evaluate intrinsic network states, building on prior evidence of baseline disruptions in this model [17]. This strategy is consistent with a broader objective in translational neuroscience: enhancing the detection of network anomalies under resting-state conditions. To deepen this analysis, we decomposed LFP spectrograms, recognizing that specific spectral components may reflect underlying cognitive and pathological states. In line with EEG based approaches [10, 33, 34] this analysis revealed functional signatures potentially tied to local cytoarchitecture. Notably, we observed patterns consistent with previously described neuron-type-specific signal features [35]. By mapping activity classifications using our proposed metric *ϵ*, we characterized spatial heterogeneity in functional groups, distinguishing disorganized (Dyslaminated) from more strucured (Eulaminated) cortical regions. We further explored the aperiodic exponent of LFP signals to asses its relationship with these functional classifications. Interestingly, a bimodal distribution emerged, with higher divergence in Dyslaminated regions and greater uniformity in Eulaminated areas. This dual pattern may reflect distinct (E/I) regimes, as previously had been suggested [36, 37], echoing previous findings from both, cognitive and epileptic contexts [7–9, 24, 25]. Under Mg^2+^-Free perturbation, this distinction faded, suggesting that strong ionic shifts may transiently override intrinsic network heterogeneity. After recovery, the absence of the bimodal pattern implies a homogenization of network activity, masking the functional distinctions evident during spontaneous states, distinctions likely rooted in cytoarchitectural organization.

These observations, together with the observed differences in inferred information transmission, support the notion that E/I balance is altered at baseline and that signal propagation may be modulated by the degree of cortical delamination, especially when exposed to external perturbation such as Mg^2+^-Free medium.

Although the original aim of this study was to characterize the network’s functional response to external perturbation, our analysis revealed that the most informative features of electrode categorization, signal aperiodicity, and intercategory transfer entropy consistently emerged from spontaneous activity. This underscores the relevance of intrinsic dynamics for uncovering structural-functional relationships and supports the broader applicability of our methodology. We propose that this methodology may be adapted to study functional inhomogeneities across diverse experimental paradigms, including healthy brain states and various neurological conditions, potentially offering a simplified approach to pre-surgical evaluation.

### 3.1 Limitations of this study

This study presents several limitations inherent to the use of an ex vivo model of cortical malformation. A primary constraint is the inability to directly validate the functional findings with histological analysis of the exact recorded regions. This is due to the use of multielectrode array (MEA) recordings in 400 *µ*m-thick cortical slices, which often sustain substantial structural damage after prolonged recording sessions, rendering subsequent histological processing unfeasible. Furthermore, while the carmustine model effectively replicates key cytoarchitectural features of focal cortical dysplasia, it does not fully capture the complexity and heterogeneity observed in human cases. Therefore, future studies applying analytical frameworks similar to human tissue, particularly samples obtained from epilepsy surgeries, will be essential to strengthen translational relevance and to assess the generalizability of the present findings.

## 4 Methods

### 4.1 Animals

Protocols followed in this study were approved by the ethics committee of the Institute of Neurobiology, UNAM (Protocol 111-A) and carried out in accordance with federal regulatory laws for animal experimentation (NOM-062-ZOO-1999). All methods were compliant with the ARRIVE guidelines and regulations. Pregnant Sprague-Dawley rats were intraperitoneally injected with BCNU following previously reported methods[12, 17]. For the Eulaminated group, *n* = 2 rats were injected with saline solution, and for the Dyslaminated group, *n* = 3 were used. All animals were kept in a 12-hour light/dark cycle room with unlimited access to food and water. A total of 5 Eulaminated and 7 Dyslaminated males were used at P30.

### 4.2 Slice Preparation

To conduct Multielectrode Array (MEA) recordings, we obtained 400 *µ*m thick coronal slices of the M1 cortex (Bregma 9.20 mm) from male rats. Rats were anesthetized with pentobarbital (60 mg/kg) and transcardially perfused with a cold sucrose-based solution[38, 39]. The brain was then removed and placed in ice-cold carbogenated solution (95% O_2_/5% CO_2_). Coronal slices were cut using a vibratome (HM-650-V, Thermo Fisher, or VT1000S, Leica) and transferred to a neuroprotective solution at 34^°^C for at least 90 minutes before use. Each recording corresponds to a different animal.

### 4.3 Electrophysiological Recordings

Recordings were obtained using a 60-electrode MEA (8 × 8 grid, 60pMEA200/30iR-Ti, Multichannel Systems), perfused from above and below with circulating ACSF at 34^°^C, bubbled with carbogen. Electrodes targeted the primary motor cortex. Negative pressure was applied to ensure adhesion of the slice. Recordings were acquired at 25 kHz for 15 minutes. Baseline activity was recorded, followed by Mg^2+^-Free ACSF recordings (3 samples), and then recovery recordings in normal ACSF (2 samples).

### 4.4 Data Analysis

#### 4.4.1 Preprocessing of Electrophysiological Data

Raw MEA recordings were decompressed and extracted from MC Rack software using McsPyDataTools^1^. We then apply a low-pass filter, using a 5th - order Butterworth filter and with a cutoff frequency of 100 Hz. Electrode traces were downsampled to 1 kHz. The noise from the power line was eliminated using a Notch Filter 37 using a quality factor of 90 We then reorganize the index of each electrode to recreate the original architecture of the MEA 60-electrode MEA (8×8 grid), and we keep the reference electrode off to continue further analysis. Quality control was performed by the comparison of the logarithmic spectrum and also revisited during the Dimension reduction of LFP. *n* = 3, Dyslaminated: *n* = 6.

#### 4.4.2 LFP Spectrograms

Spectrograms were computed using 1-second sliding windows with 75% overlap. Power spectra over 0.5–100 Hz were computed using 6 tapers and multi-taper decomposition. The number of frequency points was reduced to 60, and power was normalized using baseline mean power.

#### 4.4.3 Dimensionality Reduction and Electrode Clustering

To characterize how the spectrogram along electrodes varies for different Frequencies *f* as a function of time *t* we perform PCA in the spectrogram computed as the method described before. The number of significant dimensions goes from as many as the length of every frequency, 60 bins from 0.5-100 Hz. PCA was performed from the covariance matrix of every electrode spectrogram over the time windows used to construct it *t* and every frequency *f* as follow:

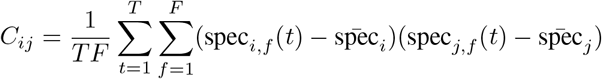

Where *T* is the number of time windows in the period considered, *F* is the number of different frequencies evaluated in each electrode. Spec_*i,f*_ (*t*) is the part of the spectrogram of electrode *i*, at that interval time in that frequency. Spec_*i*_ is the spectrogram of electrode *i*, and so on.

#### 4.4.4 Level of disarrangement metric (*ϵ*)

We took the modules that carry the largest part of the variance and then projected them into the original coordinates in the space of the total electrodes. We then performed unsupervised learning to cluster the electrodes that covariate in a similar way, using the mean shift clustering of `Sklearn` python dependency [40], with an estimated bandwidth function, using a quantile = 0.45, and a number of samples = 80.

To approximate describing the level of disarray in every MEA array, we use the projection of electrode clustering result and calculate the level of sparse of those in the original array using the following expression:

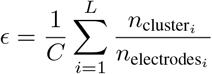

Where *C* is the total number of clusters found, *L* is the number of lines in our arrays 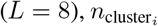 is the number of clusters that cohabit on the same line *i*, and 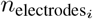 corresponds to the total number of electrodes of every line *i*, which could go from 6 to 8 due to the architecture of the array.

#### 4.4.5 Common band determination

Inspired by methodologies from Zhang, W [30], we were interested in describing a common oscillatory band along the calculated spectrograms, using mainly basal activity, and verifying if that oscillation persists in condition activities such as Mg^2+^-Free and Recovery. For this we used our normalized spectrograms and PCA was performed from the covariance matrix of each time window over the total number of electrodes (*e*) and each frequency (*f*) to construct it as follows:

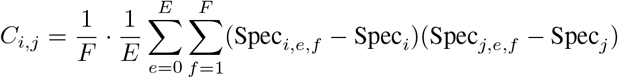

Where *F* is the number of total frequencies evaluated in the period considered, *E* is the number of different electrodes in our arrays. Spec_*i,e*_(*f*) is the part of the spectrogram at time window *i*, in an electrode *e*. Spec_*i*_ in this case represents the spectrogram in the time window *i*, etc.

We project the 4 principal components to evaluate the variation carried within as shown in Supplementary 2, and by calculating the meaning over the extreme examples (Dyslaminated, Eulaminated) in baseline activity we detected the range where the first component varies the most, to evaluate then how that range of frequencies could remain in the other records and condition activities.

#### 4.4.6 Aperiodic exponent fitting, and bimodality

Based and following the literature [9], we evaluate the aperiodic exponent in every electrode at baseline activity, Mg^2+^-Free and recovery by fitting the Lorentzian function:

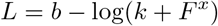

Where *b* is the broadband offset, *x* is the exponent and *k* the knee parameter, that according to Donoghue, T.and Miller, K.[9, 41] and also applied by [36] (Gao, R 2020), allows a control of the bend in the aperiodic component. *F* corresponds to the frequency vector in input frequencies. Knowing that when *k* = 0 the Lorentzian fits a line in log-log space.

To perform the tendency of the power log, we applied a Gaussian filter using `Scipy` [42] with a *σ* = 5 and also measured the variance respecting the logarithmic scale to approach the level of variation of spectrograms along all our electrodes shown in Figures 1,2.

By knowing the bimodality shape of the values at Dyslaminated and Eulaminated, as shown in Figure 2, we do both, to establish a threshold value given by our aperiodic values guiding us to the depression point that separate both density distributions, *x* = 0.6, and project the distribution of values highest and lowest to it in our real coordinates of MEA electrodes, and knowing the density variation of higher and lower values we compute the level of bimodality in our histograms following the bimodal coefficient statistic described in [43]. Finally, we applied a common linear regression between bimodality value and the *ϵ* value in baseline activity condition.

#### 4.4.7 Transfer entropy along electrode clusters

We computed the transfer entropy from baseline and also during Mg^2+^-Free and recovery conditions the common LFP band found in every cluster, ie *X*_*t*_ = LFP from cluster *n* and *Y*_*t*_ = LFP from cluster *m*; using a finite set of lags (200, (−100,100))) *L* = {*τ*_1_, *τ*_2_, …, *τ*_*k*_}. For each lag, we define the lagged version of the time series:

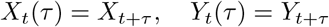

Then we calculate the Mutual Information (MI) between *X*_*t*_(*τ*) and *Y*_*t*_ as:

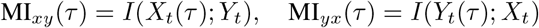

Where *I* corresponds to Shannon mutual information, and *H* to Shannon entropy:

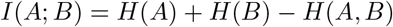

Where *A* and *B* represent the different cluster sets. To compute directionality, we took the vector MI_*xy*_(*τ*) for all *τ* ∈ *L*, where we consider the best delay as the one that maximizes the mutual information:

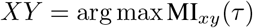

And we follow the same logic in a similar way for the other direction *Y X*. As long as we found more than two electrode clusters, we use all the possible combinations between them, to perform all possible directions.

#### 4.4.8 Permutation based statistical test

To test whether the observed mutual information was significantly greater than chance, we use a null distribution built by permutation for each lag. We generate *N* = 1000 permutations, 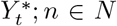 by shuffling the time series *Y*_*t*_, and vice versa for *X*_*t*_. Then we compute the equivalent MI function for every shuffle and direction. We kept with values where *p*(*τ*) *<* 0.05.

#### 4.4.9 MI of LFP components (magnitude, phase and instant frequency) during time evolution

Following the assumption the information carried in a LFP signal is distributed differently as Kayser, et al. [32] has described. We decompose our LFP in the common band evaluated (0.5–30 Hz) using a discrete Hilbert Transform [44], to obtain the phase, envelope (energy) and instantaneous frequency of every electrode.

Once having each component we computed the MI between every electrode, using a sliding window of 15 s with a 75% overlapping, using a similar expression above described:

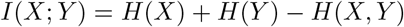

Where in this case *X* and *Y* represent the interaction between two different electrodes along MEA. We took for every electrode the maximum value of information carried along all their interactions, and we computed the same for every condition activity: Baseline, Mg^2+^-Free (3 samples), Recovery (2 samples).

This metric does not inform any directionality or causation, here we employed it as a referendum of information amount carried in every condition activity.

#### 4.4.10 Pairwise dissimilarity evaluation and Multidimensional scaling

We compute temporal dissimilarity of matrices *D*_*j*_, previously obtained due to MI temporal sliding windowing of every component of LFP signal; where each entry 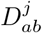 quantifies the Euclidean distance between the *a*-th entity at time *j* and the *b*-th entity at time *j* + 1, effectively decomposing the system’s temporal evolution into interpretable pairwise dissimilarities, as the following expression resumes:

Let:

*x*_*aj*_, the *a*-th row sample from *M*_*j*_

*x*_*bj*+1_, the *b*-th row sample from *M*_*j*+1_

For all *a* = 1, …, *m* and *b* = 1, …, *n*:

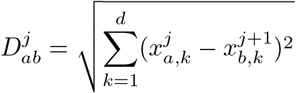

Where *k* represents the feature/dimensions index, in each sample vector. Then, with reduced vectors obtained in 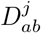, we applied MDS to assess how different the activity was from baseline, Mg^2+^-Free, to recovery. We projected the first two dimensions of the MDS and fitted an ellipse in 2D space to find the dispersion of the vectors along the length, and a Student’s t-test was performed to evaluate the difference between conditions and components.

## 5 Resource availability

The source code for processed data and all methods here reported is available on GitHub: <u>CorticalDelam-MEA</u>. The MEA recordings data can be accessed through the following DOI: <u>10.5281/zenodo.15592455</u>.

## 6 Funding

This work was financially supported by UNAM-DGAPA PAPIIT (IN203825 RRP, IN204720 LC, IA200621 HLM, IN208725 POF) and CONACYT (FC1782 LC), CONAHCYT (CF-2023-I-218 LC). Ana Aquiles received financial support from CONACYT (No. 756903).

7 Acknowledgments

We extend our sincere gratitude to the members of the Rossi-Pool Laboratory for their dedication and contributions throughout the development of this project. We are also grateful to Sergio Parra and Jerónimo Zizumbo for their valuable suggestions and feedback during the early stages of this manuscript. Special thanks go to Alessio Franci and Lucas Bayones for their insightful comments during the editing process. We acknowledge Leopoldo Gonzalez Santos for his technical support with cluster computing and Benito Ordaz, as well as the Biotherium facility at the Institute of Neurobiology. We appreciate the support of the PhD Program in Biomedical Sciences at the National Autonomous University of Mexico (UNAM).

## 8 Author contributions

A.A., RR.P., L.C., LM.H. and PO.F. contributed to the conception of the study and offered experimental method guidance. PV.L. performed the experimental procedure of MEA recordings. A.A and RR.P. designed the algorithm, performed formal analysis and visualization, and drafted the manuscript. L.C.and PO.F. provided valuable suggestions for the experiments and manuscript. All authors contributed to the review and editing of the manuscript.

## Supplementary Information

**Figure S1:**
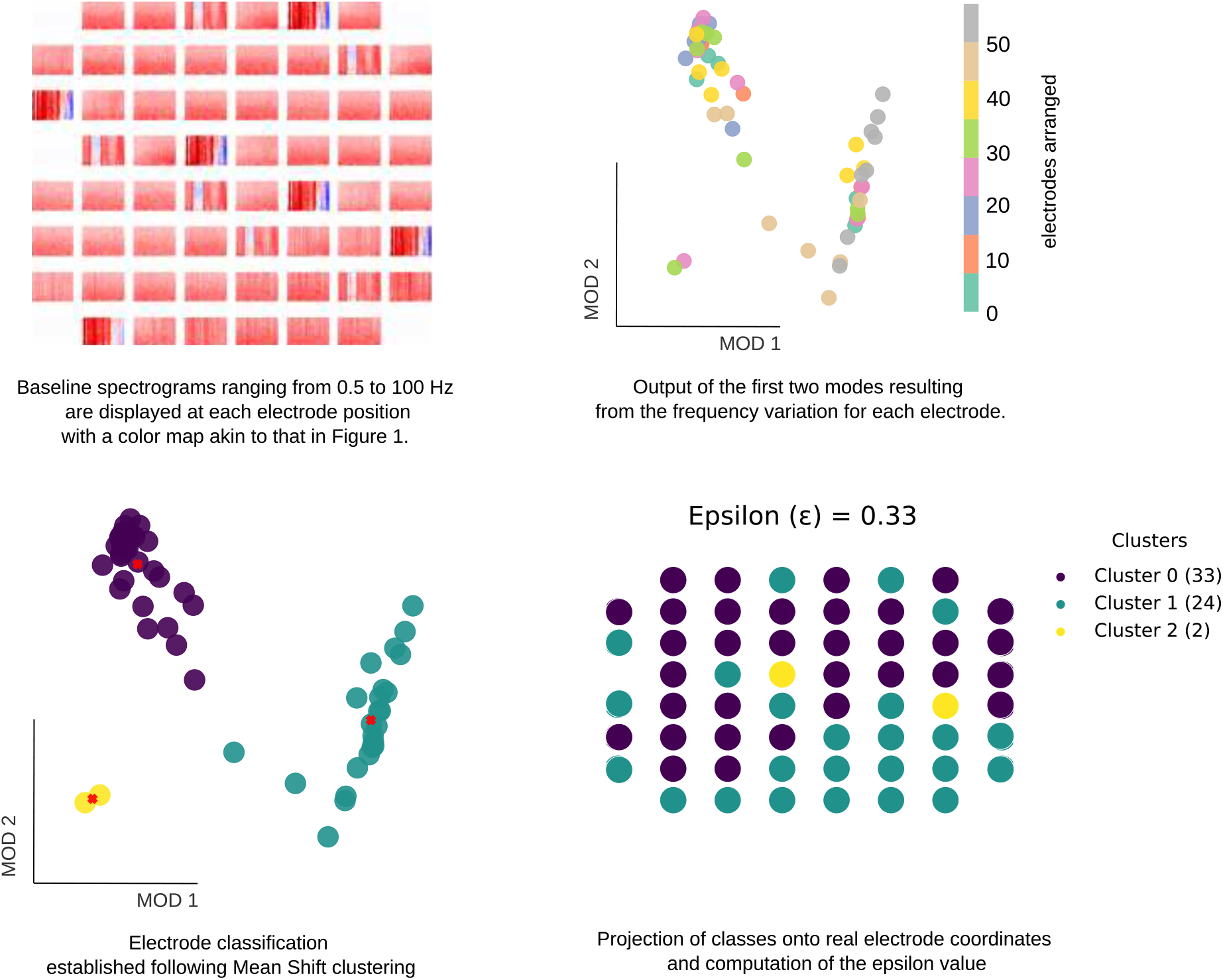
Supplementary. Resume of electrode classification method. We represent the main four steps in the computation of electrode clustering, where the source of this methodology is the baseline or spontaneous activity spectrograms, which are then covariated in the frequency domain at every electrode to identify possible electrode classes and project them into the real MEA space or real electrode coordinates, where the epsilon metric is computed. See Figure 3 and Supplementary 2.

**Figure S2:**
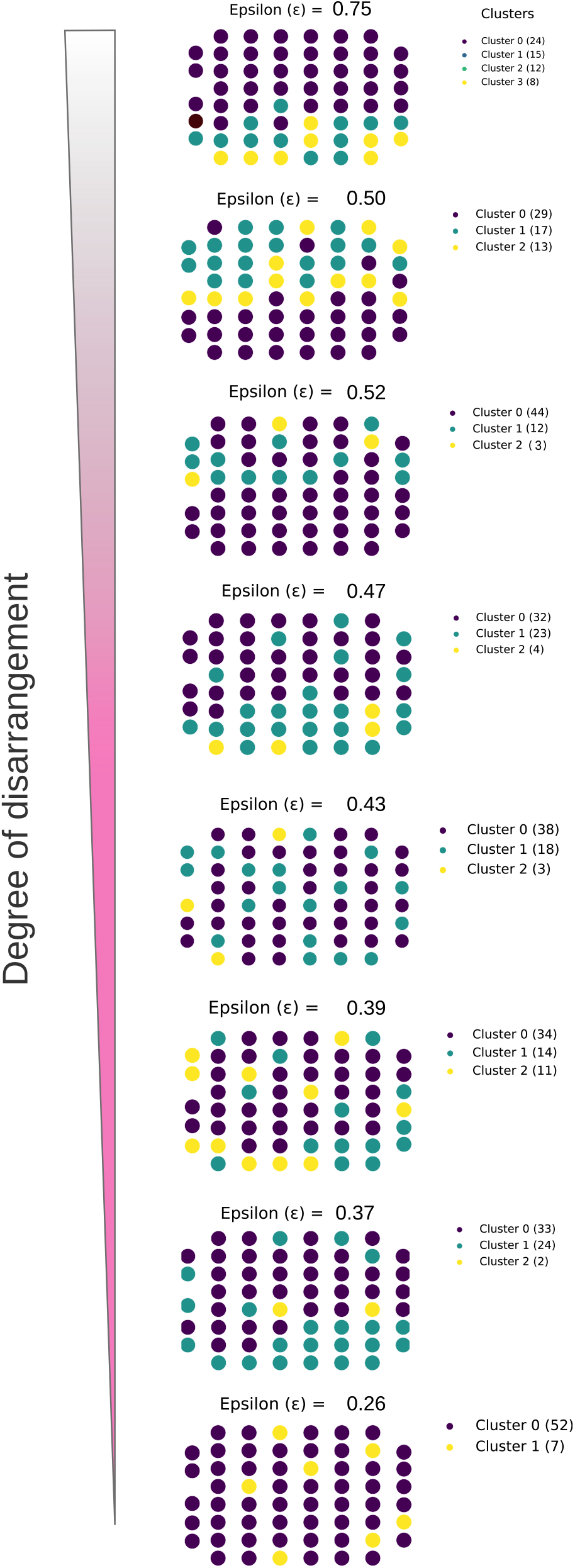
Supplementary. Degree of disarrangement in all MEA recordings. Each electrode cluster is projected into the real array position with its corresponding epsilon value and cluster description. The degradation scale shows the level of disarrangement increasing from high (pink color) to more structures (grey color).

**Figure S3:**
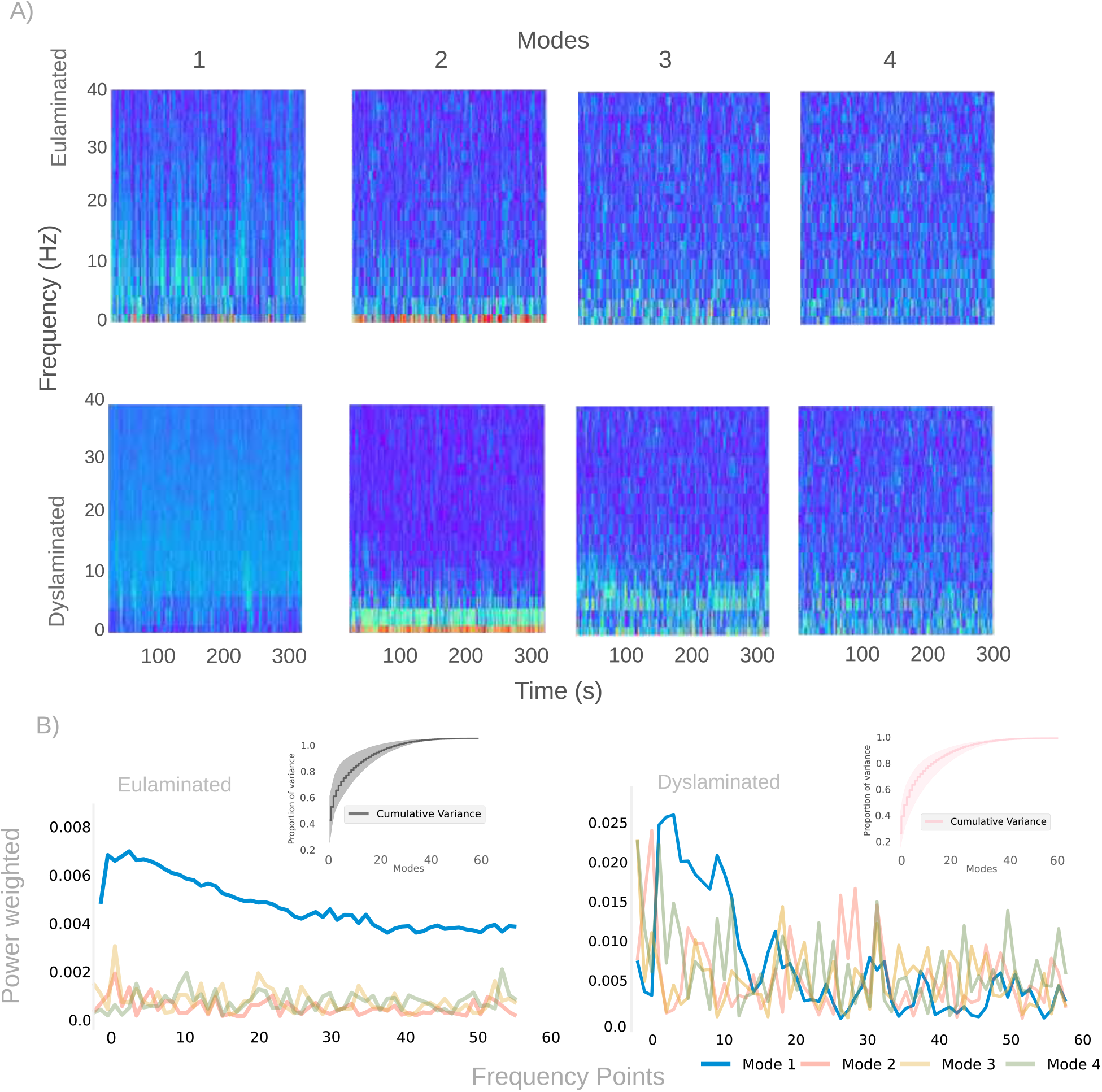
Supplementary. Common band exploration. A) Spectrogram weighted at different modes (1-4) in eulaminated, top, and dyslaminated, bottom. Each mode is representative of five minutes of baseline oscillation. B) The initial four modes have been projected onto a power spectrum. The mean cumulative variance of all the recordings has been evaluated and displayed in the top right panel, with the variance between them shown in shadow. In this context, grey and pink represent the concepts of “eulaminated” and “dyslaminated”, respectively.

## Footnotes

1 https://github.com/multichannelsystems/McsPyDataTools

